# IRF7 from the black flying fox induces a STAT1-independent ISG response in unstimulated cell lines that protects against diverse RNA viruses

**DOI:** 10.1101/2024.05.02.592239

**Authors:** Pamela C. De La Cruz-Rivera, Jennifer L. Eitson, John W. Schoggins

**Affiliations:** Department of Microbiology, UT Southwestern Medical Center, Dallas TX 75390

## Abstract

Bats are considered unique in their ability to harbor large numbers of viruses and serve as reservoirs for zoonotic viruses that have the potential to spill over into humans. However, these animals appear relatively resistant to the pathogenic effects of many viruses. Mounting evidence suggests that bats may tolerate viral infections due to unique immune features. These include evolutionary innovations in inflammatory pathways and in the molecules involved in viral sensing, interferon induction, and downstream interferon-induced antiviral effectors. We sought to determine whether interferon-stimulated genes (ISGs) from the black flying fox (*Pteropus alecto*) encoded proteins with unique antiviral activity relative to their human orthologs. Accordingly, we compared the antiviral activity of over 50 ISG human-bat ortholog pairs to identify differences in individual effector functions. We identified IRF7 from *Pteropus alecto* (Pa.IRF7) as a potent and broad-acting antiviral molecule that provides robust antiviral protection without prior activation. We show that Pa.IRF7 uniquely induces a subset of protective ISGs independent of canonical IFN signaling, which leads to protection from alphaviruses, a flavivirus, a rhabdovirus, and a paramyxovirus. In uninfected cells, Pa.IRF7 partially localizes to the nucleus and can directly bind interferon-sensitive regulatory elements (ISREs). Compared to human IRF7, Pa.IRF7 also has additional serines in its C terminal domain that contribute to antiviral activity and may serve as unique phosphorylation hubs for activation. These properties constitute major differences between bat and human IRF7 that offer additional insight into the potential uniqueness of the black flying fox immune system.

## Introduction

Bats are mammals in the order Chiroptera, which has approximately 1400 species and is the second most diverse mammalian order next to Rodentia. Targeted studies on specific viruses suggest that certain bat species are carriers of several viruses that are pathogenic in humans, including henipaviruses, coronaviruses, and filoviruses (Chua et al., 2002; Halpin et al., 2000; W. Li et al., 2005; Towner et al., 2009). Recent studies using broad analytical methods further indicate that the order Chiroptera has a particularly high richness of virulent zoonotic viruses (Gibb et al., 2020; Guth et al., 2022; Mollentze & Streicker, 2020; Olival et al., 2017). These studies have shown that, in addition to the viruses noted above, bats also harbor diverse viral groups such as flaviviruses, arenaviruses, and rhabdoviruses. The ability of bats to harbor large numbers of viruses has led to considerable interest in these animals with respect to virus-host interactions and antiviral immunity.

The antiviral innate immune response is highly conserved in mammals, and bats are no exception with respect to the key molecules that mediate this response. In mammals, viruses activate the innate immune response by triggering cellular sensors that usually recognize viral nucleic acid. The sensors activate a signal transduction cascade whereby interferon regulatory factors (IRFs) transcriptionally induce interferon (IFN) gene expression. IFNs are translated, secreted from the cells, and function as cytokines to elicit a broad transcriptional program mediated by the JAK/STAT signaling pathway. This culminates in the induction of hundreds of interferon-stimulated genes (ISGs), many of which encode effector proteins with antiviral activity (Ivashkiv & Donlin, 2014; Platanias, 2005; Schoggins, 2019).

Despite the highly conserved nature of this pathway in mammals, recent studies suggest that specific aspects of immunity may be unique in bats. With respect to mechanisms that may allow viruses to tolerate infection, certain bat species have been shown to have a functionally impaired NLRP3 inflammasomes or a complete loss of inflammatory PHYIN genes (Ahn et al., 2019; Ahn et al., 2016). With respect to antiviral pathways, relative to humans, some bats have greater numbers of IFN genes, while other bats have a contracted IFN gene repertoire (Demian et al., 2024). The biological relevance of IFN gene numbers in bats, or any other IFN-expressing vertebrate, is not known. Some bats have dampened STING activity, which affects downstream IFN induction (Xie et al., 2018). IFN signaling, while largely conserved in bat cells, appears to have distinct temporal kinetics of ISG induction when compared to analogous signaling in human cells (De La Cruz-Rivera et al., 2017).

Although great efforts have been made to expand our knowledge of the transcriptional signatures in response to IFN or viral infection in a number of different bat species (Arnold et al., 2018; Biesold et al., 2011; De La Cruz-Rivera et al., 2017; Gamage et al., 2022; Glennon et al., 2015; Holzer et al., 2016; Holzer et al., 2019; Pasquesi et al., 2022; Pavlovich et al., 2020; Wu et al., 2013; Wynne et al., 2014; Wynne et al., 2017; Zhang et al., 2017), studies examining the function of bat antiviral effectors are very limited. In a previous study, we screened an IFN-induced cDNA library from the black flying fox and found that the ISG RTP4 potently inhibited flaviviruses (Boys et al., 2020). Here, we used a distinct approach to identify antiviral genes in bats. We generated a relatively small bat ISG expression library based on homology differences between human and bat ISGs. We overexpressed select bat ISGs and their human orthologs and compared their ability to inhibit viral infection. We found that compared to human IRF7 (*Homo sapiens*, Hs.IRF7), bat IRF7 (*Pteropus alecto*, Pa.IRF7) had significantly increased antiviral capacity in cells lacking canonical JAK-STAT signaling. This augmented antiviral activity was due to Pa.IRF7 directly regulating the transcription of numerous ISGs that protected cells from several diverse RNA viruses. In mechanistic studies, we found that Pa.IRF7 readily localizes to the nucleus in the absence of stimulation, directly binds to ISREs in the promoters of ISGs, and contains a C terminal domain harboring functionally relevant serines that are not present in Hs.IRF7.

## Results

### Bat-Human ISG Ortholog Antiviral Screen

To compare the antiviral activities of human and bat ISG orthologs, we first established criteria for inclusion of genes in a mini-ISG library. We focused primarily on the black flying fox (species *Pteropus alecto*), a known viral reservoir whose genome has been sequenced (Zhang et al., 2013) and characterized at the level of IFN responsiveness (De La Cruz-Rivera et al., 2017; Janardhana et al., 2012; Virtue et al., 2011; Zhou et al., 2013; Zhou et al., 2016). Our previously annotated list of approximately 450 human ISGs (Schoggins et al., 2011) was submitted to Batch Entrez to obtain protein sequences in FASTA format. The protein sequences were submitted to NCBI BLAST using *P. alecto* as the query genome. The retrieved bat sequences were compared to their human orthologs, and proteins with greater than 70% identity at the amino acid level, which we hypothesized may have similar functions, were removed. We additionally removed all genes greater than 4000 base pairs to reduce difficulties in cloning, as well as ISGs encoding immunomodulatory or secreted proteins, such as major histocompatibility (MHC) proteins and chemokines. Several human ISGs were not adequately annotated in the *P. alecto* genome, and their orthologs were obtained from *Pteropus vampyrus*. The final bat ISG list contained 78 ISGs, which were codon optimized, synthesized, and cloned into Gateway compatible lentiviral vector plasmids (**Table S1**). Several human orthologs of bat genes were not available in our ISG cDNA library, leaving approximately 69 human-bat ISG ortholog pairs.

To identify differences in antiviral function of ISGs between bat and human orthologs, we used our previously described flow cytometry-based screening assay (Schoggins et al., 2011) (**Fig 1A**). Briefly, lentiviral vectors co-expressing individual ISGs or a firefly luciferase (Fluc) control and a red fluorescent protein TagRFP were used to transduce *STAT1^−/−^* human fibroblasts. Cells were then infected with GFP-expressing reporter viruses, including the alphaviruses O’nyong nyong virus (ONNV), Sindbis virus (SINV), and Venezuelan equine encephalitis virus (VEEV), the rhabdovirus vesicular stomatitis virus (VSV) and the flavivirus yellow fever virus (YFV). The percentage of GFP+ cells was quantified by flow cytometry. Certain genes were triaged for several reasons, including poor lentivirus production, poor lentivirus transduction, or highly disparate levels of the RFP reporter between specific human-bat ISG lentiviral pairs. From the original 69-gene set, we were able to obtain reliable data for 42 human-bat ISG pairs in the VEEV screen and 50 to 55 ISG pairs for all other viruses. We calculated z-scores and plotted the human ISG z-score versus the bat ISG z-score (**Fig 1B, Table S2**). Many ISGs were close to the origin, indicating little to no effect on viral infection. ISGs with similar phenotypes between the two orthologs (−1.5 < z < 1.5) fall near the diagonal, with inhibitory and enhancing ISGs in the lower left and upper right quadrants, respectively. We observed broad antiviral activity with both bat and human IRF1, our internal control based on previous findings (Schoggins et al., 2014; Schoggins et al., 2011). For most ISGs, the bat and human orthologs had similar effects on viral infectivity. For example, both bat and human IFI6 inhibited the flavivirus yellow fever virus (YFV)(Richardson et al., 2018), and both MAP3K14 orthologs inhibited the alphaviruses (Liu et al., 2019), as previously reported. We also observed conserved phenotypes with a subset of genes previously shown to enhance viral infection, including ADAR1 and LY6E (Schoggins et al., 2011). Notably, for several genes, including IRF7 and IFIT1, the bat ortholog had increased antiviral activity relative to the human ortholog (**Fig 1B**, **Table S2**). We were intrigued by the IRF7 phenotype, as our screens were performed in *STAT1*^−/−^ cells that do not respond to IFN (Dupuis et al., 2003). Human IRF7 (Hs.IRF7) classically exhibits an antiviral effect by inducing the production of antiviral IFNs, which lead to expression of ISGs via JAK-STAT signaling (Au et al., 1998; Lin, Mamane, et al., 2000; Marie et al., 1998; Sato et al., 2000). Hs.IRF7 had minimal antiviral effects in *STAT1*^−/−^ cells, as predicted, whereas Pa.IRF7 robustly suppressed infection. This suggested that Pa.IRF7 may have unique STAT1-independent antiviral properties.

**Figure 1:**
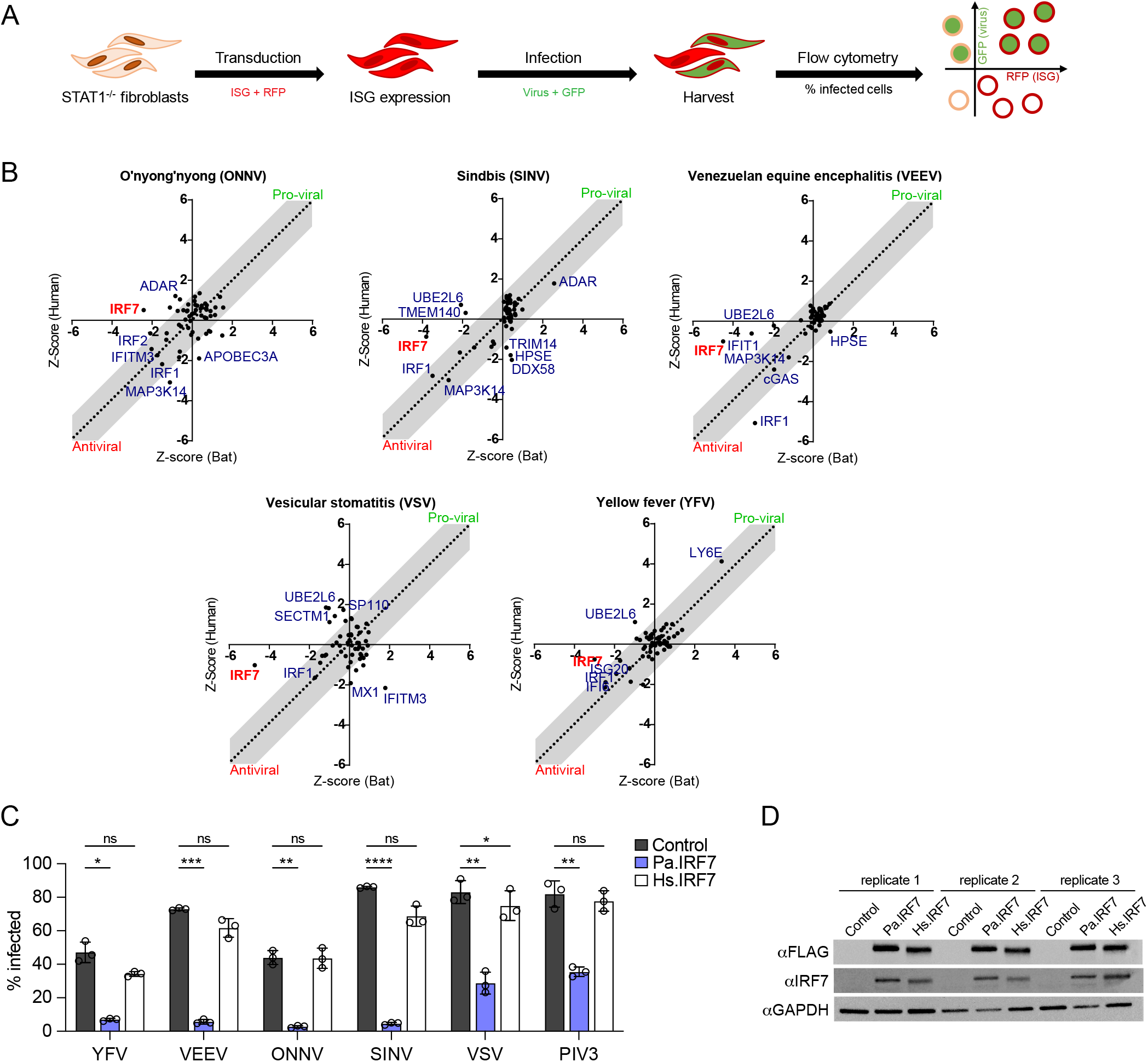
Pa.IRF7 is a broadly-acting antiviral ISG in human *STAT1*^−/−^ fibroblasts. (A) Human *STAT1*^−/−^ fibroblasts were transiently transduced with a lentiviral vector encoding a single human or bat ISG and RFP reporter. Cells were then infected with a GFP-expressing virus and infection was quantified using flow cytometry. Each screen was performed with two independent replicates (B) Z-score plots for all screens. ISGs that inhibit viral infection have negative z-scores and are found on the bottom left. ISGs that promote viral infection have positive z-scores and are in the upper right. Orthologs with similar activity are located inside of the grey bars, indicating ortholog z-scores are within 1.5 of each other. Data represent the average z-score of duplicate screens. (C) *STAT1*^−/−^ fibroblasts were transduced with lentivirus to stably express FLAG-tagged Pa.IRF7 or Hs.IRF7, or vector control and then selected with puromycin. Stable cell lines were infected with the indicated viruses at an MOI of 1.5 (YFV), 4 (VEEV), 2 (ONNV), 12 (SINV), 30 (VSV), and 15 (PIV3) for the following time periods: YFV (24h), VEEV (5h), ONNV (18h), SINV (18h), VSV (4h), and PIV3 (18h). Infectivity was quantified by flow cytometry and is presented as means +/− SD, n=3. Statistical significance was determined by one-way ANOVA with Dunnett’s multiple comparison’s test. *, P<0.05; **, P<0.01; ***, P<0.001; ****, P<0.0001, ns, not significant. (D) Matched western blots from experiments above showing similar expression levels of FLAG-tagged IRF7s, n=3.

We sought to corroborate these findings and obtain additional mechanistic insight to understand potential unique properties of IRF7. To confirm the results from our screen, we used drug-selectable lentiviral vectors to make *STAT1^−/−^* cell lines that stably overexpress an empty vector control, Pa.IRF7, or Hs.IRF7. We infected the cells with the five viruses from the original screens and the paramyxovirus parainfluenza virus type 3 (PIV3), which was chosen because pteropid bats are considered reservoirs for related henipaviruses in the *Paramyxoviridae* family. For these validation assays, we also changed the coding sequence of Pa.IRF7. The original coding sequence obtained from the database was based on an accession number (ELK13169.1) that lacked 30 amino acids relative to an updated, full-length Pa.IRF7 accession number (XP_006911017.1). Similar to our initial screen, in these validation assays, Pa.IRF7 was broadly antiviral while Hs.IRF7 had little to no antiviral activity in these cells (**Fig 1C**). The differential antiviral activities were not due to differences in protein abundance, as both proteins were expressed similarly (**Fig 1D**). Summarily, this novel ISG ortholog screening strategy led to the identification of Pa.IRF7 as having significantly greater antiviral effects than Hs.IRF7.

### Pa.IRF7 does not require type I IFN signaling for antiviral activity

Next, we tested IRF7 phenotypes in HeLa cells, which have intact interferon induction and signaling pathways (K. Li et al., 2005; Vitali & Scadden, 2010; Yoneyama et al., 2004). *STAT1^−/−^* fibroblasts and HeLa cells stably expressing an empty vector control, Pa.IRF7, or Hs.IRF7 were infected with YFV and SINV. In *STAT1^−/−^* fibroblasts, only Pa.IRF7 inhibited infection, whereas both Hs.IRF7 and Pa.IRF7 potently inhibited YFV and SINV in HeLa cells (**Fig 2A, 2B**). The phenotype in HeLa cells is likely due to IRF7-mediated induction of IFN, as shown in previous reports (Au et al., 1998; Lin, Genin, et al., 2000; Lin, Mamane, et al., 2000; Marie et al., 1998; Sato et al., 2000). Western blot analysis of cell lysates indicated that STAT1 was undetectable in *STAT1^−/−^* fibroblasts but present and inducible by IRF7 expression in HeLa cells (**Fig 2C**). In addition, a downstream ISG effector, IFIT1, was induced by both Pa.IRF7 and Hs.IRF7 in HeLa cells. Notably, despite defective IFN signaling in *STAT1^−/−^* fibroblasts, expression of Pa.IRF7, but not Hs.IRF7, resulted in induction of IFIT1 (**Fig 2C**).

**Figure 2:**
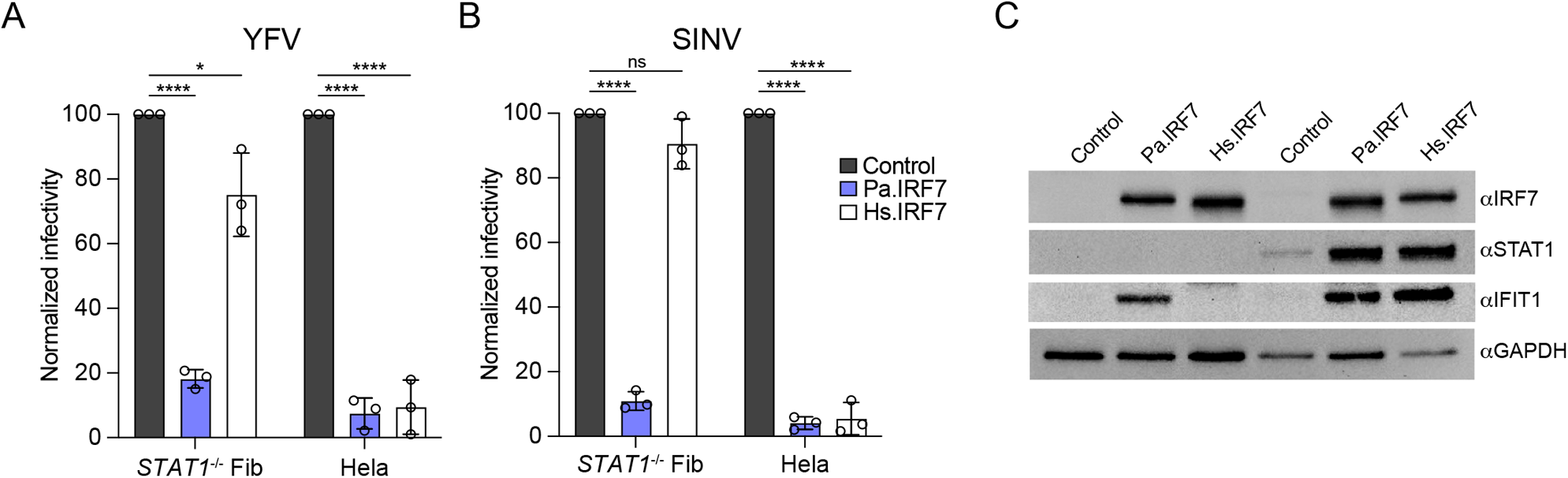
Bat IRF7 does not require type I IFN signaling for antiviral activity. (A,B) *STAT1*^−/−^ fibroblasts (A) or HeLa cells (B) were transduced with lentivirus to stably express FLAG-tagged Pa.IRF7 or Hs.IRF7, or control, then infected with YFV at an MOI of 1.5 for 24h or SINV at MOI of 12 for 18h. Infectivity was quantified by cytometry and normalized to control cells. Data are presented as means +/− SD, n=3. Statistical significance was determined by one-way ANOVA on non-normalized data with Dunnett’s multiple comparison’s test. *, P<0.05; ****, P<0.0001, ns, not significant. (C) Western blots from lysates in A and B, showing abundance of IRF7, STAT1, IFIT1, and GAPDH. Data are representative of three similar experiments.

### Pa.IRF7 induces multiple ISGs

As Pa.IRF7 is a transcription factor that inhibited diverse viruses and induced IFIT1 in a STAT1-independent manner, we asked whether Pa.IRF7 expression conferred a unique transcriptional signature. We isolated total RNA from *STAT1^−/−^* fibroblasts expressing an empty vector cassette, Pa.IRF7, or Hs.IRF7 and assessed global transcriptional changes by bead-based microarray. Despite lack of any immune stimulation, cells expressing Pa.IRF7 had significantly increased abundance of over 50 genes, most of which are classified as ISGs (**Fig 3A**) (Der et al., 1998; Rusinova et al., 2013; Shaw et al., 2017). Several genes had reduced expression in IRF7-expressing cells, and the magnitude of down-regulation was similar with each ortholog (**Fig 3B**). Additionally, the expression levels of individual type I, II and III IFNs and their receptors were not significantly different between cells expressing vector control or either IRF7 ortholog (**Fig 3C**). Increased abundance of mRNAs for the ISGs *OAS2*, *IFI44L*, and *RSAD2*/viperin in *STAT1^−/−^*fibroblasts expressing Pa.IRF7 was confirmed by RT-qPCR (**Fig 3D**). In contrast to the data obtained from microarray analysis, the more sensitive RT-PCR assay revealed that *IFNB1* mRNA was modestly induced—approximately 5-fold, and to similar extents by both bat and human IRF7.

**Figure 3:**
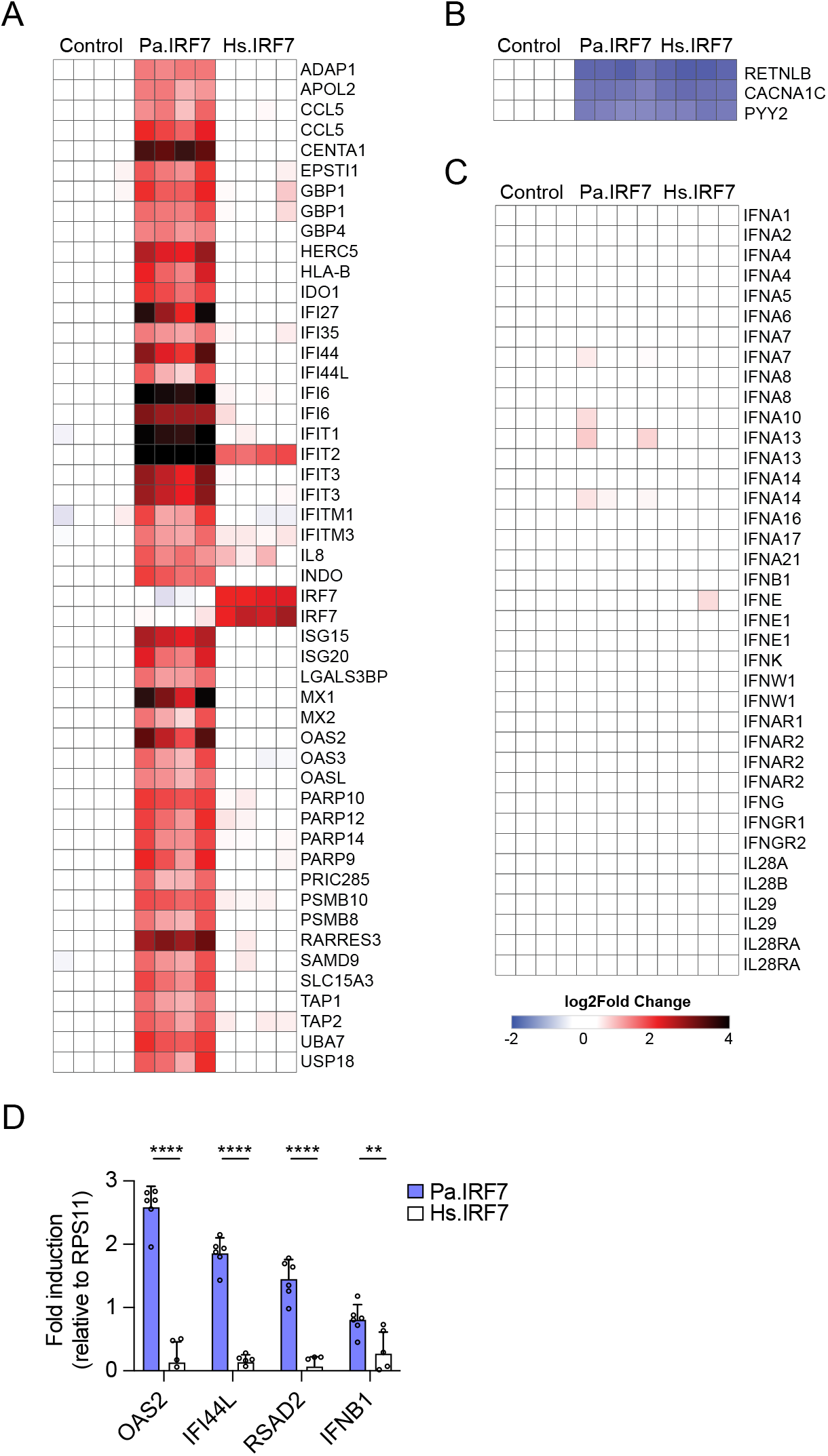
Bat IRF7 expression increases ISG mRNAs in unstimulated *STAT1*^−/−^ fibroblasts. (A-C) RNA was isolated from *STAT1*^−/−^ fibroblasts stably expressing vector control, Pa.IRF7, or Hs.IRF7 and microarray analysis was run on the Illumina Human-HT-12 V4 chip. Expression of genes that were significantly (p<0.05) upregulated (A) or downregulated (B), or expression of selected IFN and IFN receptor genes is depicted as heatmaps (C). Each column represents one of four independent experiments per cell line. (D) RT-qPCR analysis of OAS2, IFI44L, RSAD2, and IFNB1 mRNA from *STAT1*^−/−^ fibroblasts expressing vector control, Pa.IRF7, or Hs.IRF7. Relative gene expression was normalized to vector control cells using RPS11 as a reference gene. Data is presented as mean ± SD, n=6. Statistical significance was determined by t-test of log-normalized data. ***, P<0.001; ****, P<0.0001.

### Endogenous Pa.IRF7 controls viral infection and ISG induction in bat cells

Our data thus far demonstrate enhanced antiviral effects of Pa.IRF7 when it is ectopically expressed in human STAT1-deficient cells. To determine whether the expanded ISG-inducing ability of Pa.IRF7 occurs in species-matched cells, we used CRISPR-Cas9 gene editing to silence IRF7 in immortalized kidney cells from the black flying fox, herein referred to as “PaKi” cells (Crameri et al., 2009). We first generated three PaKi IRF7-deficient single-cell clones, selected for their loss of IRF7 expression upon IFN stimulation (**Fig 4A**). We then infected parental PaKi and the three clonal lines with YFV in the presence or absence of IFN stimulation. In unstimulated cells, there were no differences in YFV infectivity across the cell lines (**Fig 4B**). When cells were pre-treated with IFN, YFV was suppressed in parental PaKi cells, and this repression was partially relieved in two of the IRF7-deficient clonal cell lines (**Fig 4B**). We selected clone 4G9, herein referred to as “IRF7 KO,” for further analysis. This clone had a deletion of a cytosine at position 383 (g.383delC) of the *IRF7* gene, which was confirmed by PCR amplification and Sanger sequencing (**Fig 4C**). The deletion induces a frameshift mutation within the IRF7 coding sequence, significantly altering the amino acid sequence after the mutation site and introducing a premature stop codon at position 159. When parental PaKi or IRF7 KO cells were treated with IFN, IRF7 KO cells exhibited decreased induction of the ISGs *OAS1* and *USP18* (**Fig 4D**), suggesting that IRF7 is required for their maximal induction during an IFN response.

**Figure 4.**
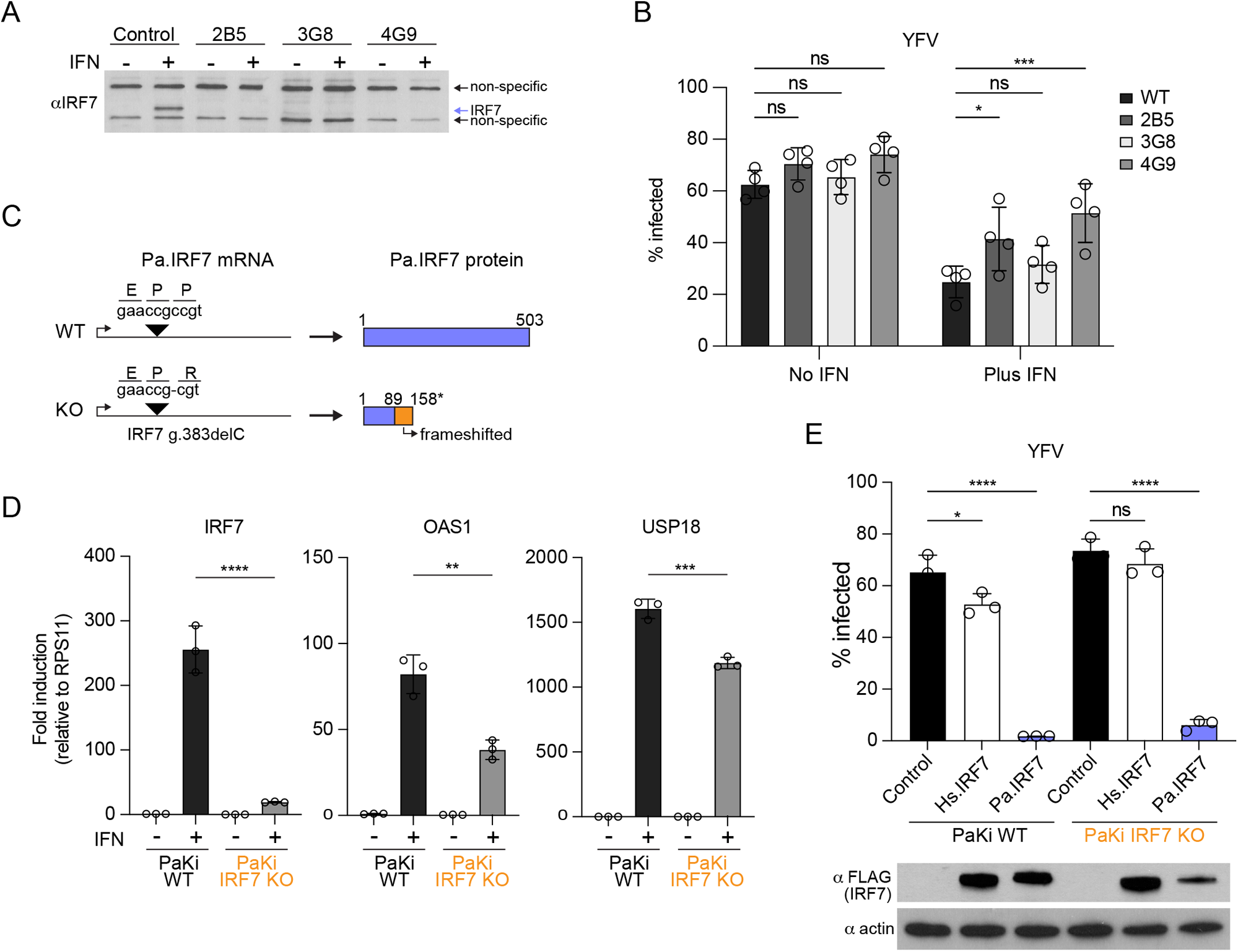
Pa.IRF7 contributes to ISG expression and antiviral activity in bat cells. (A) Lysates from PaKi WT or IRF7 KO single cell clones were assayed by Western blot for IRF7 expression in the absence and presence of 100U/mL universal IFNα for 24h. (B) WT or IRF7 KO PaKi clones were infected with YFV at an MOI of 1.5 for 24 h in the absence or presence of 100 U/mL universal IFN. Infectivity was quantified by flow cytometry and presented as means +/− SD, n=3. Statistical significance was determined by one-way ANOVA with Dunnett’s multiple comparison’s test. *, P<0.05; ***, P<0.001, ns, not significant. (C) Sequencing of genomic DNA from WT or IRF7 KO single cell clone 4G9 confirmed genetic deletions as indicated. Protein products are shown on the right. (D) RT-qPCR analysis of bat IRF7, bat OAS1, and bat USP18 PaKi or IRF7 KO cells without or with IFN treatment. Relative gene expression was normalized to vector control cells using bat RPS11 as a reference gene. Data is presented as mean ± SD, n=3. Statistical significance was determined by t-test of log-normalized data. **, P<0.01; ***, P<0.001; ****, P<0.0001. (E) (Top) PaKi or IRF7 KO cells were transduced with lentivirus to stably express Pa.IRF7 or Hs.IRF7, or control, then infected with YFV at an MOI of 1.5 for 24 h. Infectivity was quantified by cytometry and is presented as means +/− SD, n=3. Statistical significance was determined by one-way ANOVA with Dunnett’s multiple comparison’s test. *, P<0.05; ****, P<0.0001, ns, not significant. (Bottom) Representative western blot from one replicate showing expression of ectopically expressed IRF7 and actin.

To assess antiviral effects of Hs.IRF7 and Pa.IRF7 in bat cells, we transduced parental PaKi and IRF7 KO cells with lentivirus expressing both IRF7 orthologs. In PaKi cells, ectopic expression of Pa.IRF7 strongly suppressed YFV, whereas Hs.IRF7 expression only modestly reduced virus infectivity. In IRF7 KO cells reconstituted with IRF7 orthologs, Pa.IRF7 but not Hs.IRF7 inhibited YFV (**Fig 4E**). Together, these data indicate that in an orthologous cell background, IRF7 from the black flying fox contributes to ISG induction and viral inhibition.

### Pa.IRF7 readily localizes to the nucleus and directly binds ISG promoters

Next, we investigated the subcellular localization and function of Pa.IRF7 as compared to Hs.IRF7. IRF7 normally resides in the cytoplasm and translocates to the nucleus upon activation by viral infection (Sato et al., 1998). As ectopic expression could alter localization patterns, we examined endogenous IRF7 localization in human and bat cells. In human THP-1 monocytes, Hs.IRF7 was nearly undetectable in unstimulated cells, and Hs.IRF7 abundance increased with IFN treatment, as expected (**Fig 5A**). We fractionated cell lysates into cytosolic and nuclear compartments and quantified the amount of IRF7 in each fraction. Only 1% to 6% of Hs.IRF7 localized to the nucleus in IFN-treated THP1 across three experiments (**Fig 5B**).

**Figure 5.**
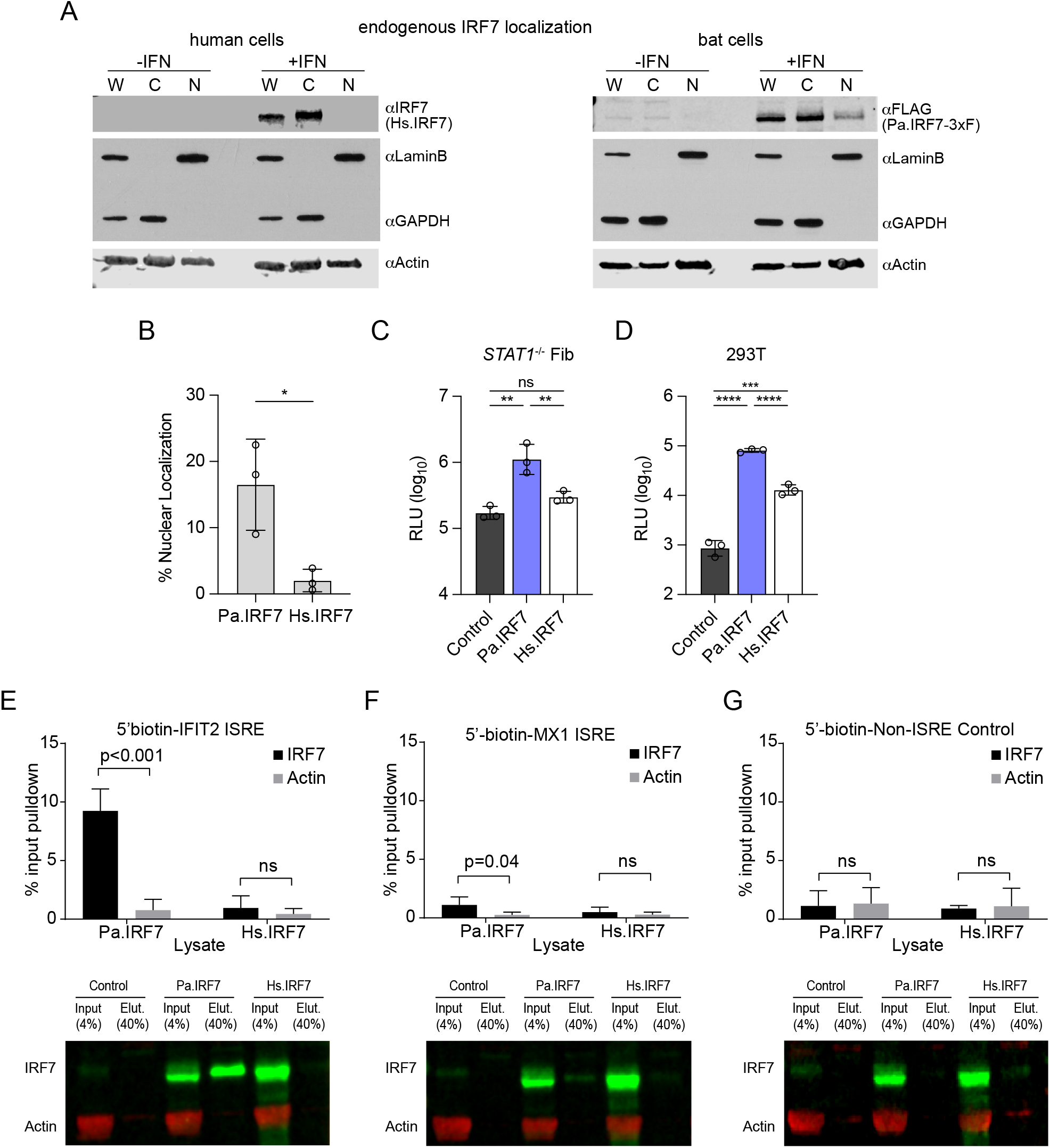
Pa.IRF7 readily localizes to nucleus and binds ISREs. (A) Human THP1 cells (left) or PaKi cells (right) bearing a 3xFLAG tag fused to the genomic IRF7 locus were treated with IFN and lysates were fractionated into nuclear and cytosolic compartments. Western blots of whole cell lysate (W), nuclear (N), and cytosolic (C) fractions were probed with antibodies: anti-IRF7 to detect Hs.IRF7, anti-FLAG to detect endogenous Pa.IRF7, anti-laminB for nuclear protein, anti-GAPDH for cytosolic protein, anti-actin for loading control. (B) Quantification of N/C ratio of data from (A), presented as mean +/− SD, n=3. Statistical significance was determined by paired t-test. *, P<0.05. (C, D) *STAT1*^−/−^ fibroblasts (C) or HEK293 (D) expressing Pa.IRF7, Hs.IRF7 or empty vector were transfected with a plasmid that expresses luciferase under control of the hamster IFIT2 ISRE and the luciferase signal was measured after 24h. Data are presented as mean +/− SD, n=3. Statistical significance was determined by one-way ANOVA with Dunnett’s multiple comparison’s test. **, P<0.01; ***, P<0.001; ****, P<0.0001, ns, not significant. (E-G) Lysates from *STAT1*^−/−^ fibroblasts stably expressing Pa.IRF7 or Hs.IRF7 were incubated with 5’-biotin-labeled DNA probes overnight. DNA-protein complexes were pulled down using streptavidin beads and the elution was probed for IRF7 and actin as a negative control. Quantification of percent input that was eluted was calculated for both IRF7 and actin for probes (E) IFIT2, (F) MX1 and (G) a sequence with no known promoter elements. Data is presented as mean +/− SD, n=3. Statistical significance was determined by t-test. *, P<0.05; ***, P<0.001; ns, not significant. A representative gel is shown under the corresponding graph.

To examine Pa.IRF7 subcellular localization in bat cells, we initially had difficulty resolving compartmentalized expression with available IRF7 antibodies. We thus used CRISPR-mediated homology directed repair to insert DNA sequences encoding a 3xFLAG tag fused to C terminus of the Pa.IRF7 genomic locus (Pa.IRF7-3xF) (**Fig S1A**). This allowed us to detect endogenous Pa.IRF7 more easily with anti-FLAG antibodies. Several single cell clones were grown and one of them, clone 4, had robust induction of Pa.IRF7-3xF when cells were treated with IFN (**Fig S1B**). A time course of IFN treatment showed rapid induction of Pa.IRF7-3xF that peaked at approximately 12 hours after treatment (**Fig S1C**). When expressed via lentivirus, the FLAG-tagged Pa.IRF7 showed comparable antiviral activity to the untagged protein (**Fig S1D**). We then examined subcellular localization of Pa.IRF7-3xF in these cells. Pa.RF7-3xF was already detectable at low levels in unstimulated cells, in contrast to undetectable Hs.IRF7 abundance in unstimulated human THP-1 cells (**Fig 5A**). In response to IFN treatment, expression of Pa.IRF7-3xF was markedly induced, and 9-22% of the protein localized to the nucleus across three experiments. This level of nuclear localization was significantly higher than that observed for Hs.IRF7 (**Fig 5B**). These data suggest that Pa.IRF7-dependent effects on ISG expression may be due to a fraction of the protein readily translocating to the nucleus in the absence of a stimulus.

Based on these data, we hypothesized that Pa.IRF7 was functioning through ISREs to induce ISG expression. To test this, we transfected a plasmid encoding a firefly luciferase gene (Fluc) driven by the IFIT2 ISRE into *STAT1^−/−^* fibroblasts stably expressing vector control or each IRF7. Cells expressing Pa.IRF7 induced higher ISRE-dependent Fluc expression than cells expressing Hs.IRF7, which did not activate the Fluc reporter beyond the control (**Fig 4C**). We performed a similar assay in 293T cells using transient transfection of the ISRE reporter plasmid and IRF7 plasmids. In these cells, which constitutively express STAT1 (Liu et al., 2001), we observed ISRE activation by both IRF7s, with Pa.IRF7 inducing the reporter at a level that was 10-fold higher than that induced by Hs.IRF7. We speculate that in 293Ts, both IRF7s induce some amount of IFN which activates the ISRE, and that Pa.IRF7 additionally activates IFN-independent expression of the ISRE, though additional studies are needed to examine this further. Together, these data suggest that expression of Pa.IRF7 leads to activation of ISREs to induce ISG expression, consistent with gene expression profiling (**Fig 3**).

Next, we tested whether Pa.IRF7 directly binds ISREs. We used 5’-biotinylated primers to amplify regions of the human genome containing ISREs from IFIT2 or MX1, two of the genes induced to high levels in *STAT1^−/−^*fibroblasts expressing Pa.IRF7 as compared to Hs.IRF7 (**Fig 3A**). The DNA probes were incubated with lysates from cells stably expressing vector control or each IRF7, and complexes were pulled down using streptavidin beads. Samples were eluted from the beads, and Western blot was used to detect IRF7 and actin as a nonspecific control (Fig 4E-G). Using the IFIT2 ISRE probe, approximately 8% of input Pa.IRF7 was pulled down, indicating direct binding of IRF7 to the IFIT2 promoter (**Fig 4E**). In contrast, the percentage of Hs.IRF7 pulled down was not significantly increased over that of actin. Using the MX1 probe, approximately 1% of input Pa.IRF7 was pulled down (**Fig 4F**). Although less Pa.IRF7 bound the MX1 promoter than the IFIT2 probe, the binding was still significant relative to actin, and Hs.IRF7 was not pulled down using the MX1 promoter. Neither IRF7 ortholog bound significantly to a non-ISRE negative control probe (**Fig 4G**). These results provide mechanistic data to support a model in which Pa.IRF7, even in unstimulated cells, is capable of directly binding ISG ISREs and causing STAT1-independent induction of antiviral ISGs.

### The C terminal serine-rich region of Pa.IRF7 is important for antiviral activity

Our studies suggest that that Pa.IRF7, unlike its human counterpart Hs.IRF7, becomes transcriptionally active without requiring a stimulus such as a viral infection. To understand the basis for this difference, we aligned the amino acid sequences of Hs.IRF7 and Pa.IRF7 and determined they were 64% identical and 70.7% homologous (**Fig S2**). This comparison revealed disparate levels of homology across the various IRF7 domains, with the constitutive activation domain (CAD) showing the most variation (**Fig 6A**). We also noted that the serine-rich C-terminal domain (CT) of Pa.IRF7 has three additional serines compared to Hs.IRF7 (12 vs. 9 serines) (**Fig 6B**). Serines are pivotal for the transcriptional activity of human and murine IRF7 and serve as kinase substrates within the C-terminal domain (Marie et al., 1998; Marie et al., 2000). Specifically, serines 477 and 479 in Hs.IRF7 are critical for type I IFN gene induction following viral infection (Lin, Mamane, et al., 2000). Phosphorylation at these sites by IKKε or TBK1 triggers nuclear translocation of IRF7 and its direct interaction with the IFNβ promoter, enhancing IFN gene transcription (Fitzgerald et al., 2003; Sharma et al., 2003). Substitution of these residues with phosphomimetic aspartic acid increases transcription of IFN genes by inducing nuclear translocation and direct binding of IRF7 to the IFNβ promoter in both infected and uninfected cells (Lin, Mamane, et al., 2000). Pa.IRF7 preserves all of the serines found in Hs.IRF7, but also has three additional series at positions corresponding to Gly464, Leu480 and Ala485, the latter two being in the vicinity of known phospho-sites Ser477/479.

**Figure 6:**
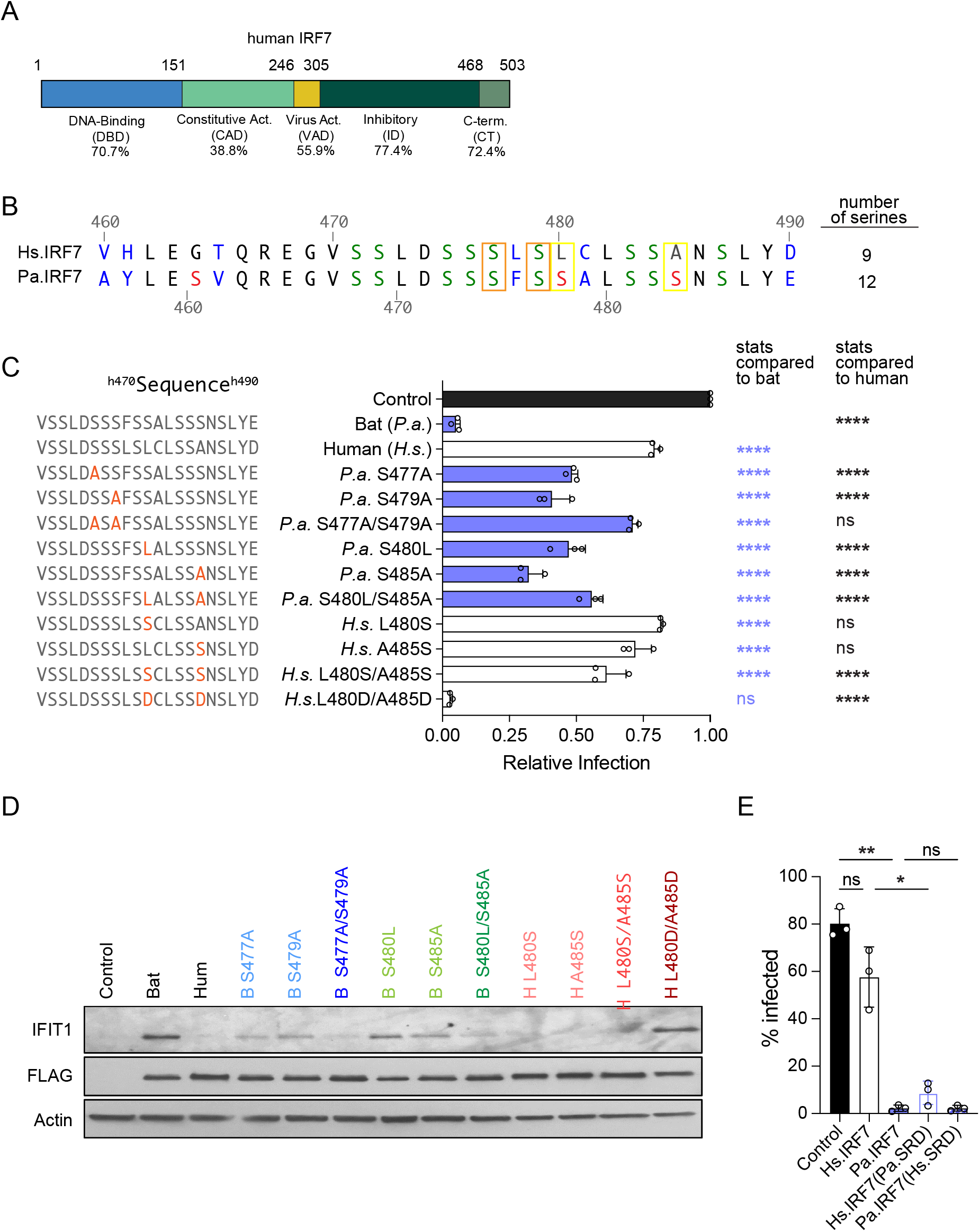
Additional serines at C terminus contribute to augmented antiviral activity of Pa.IRF7. (A) Cartoon diagram of Hs.IRF7 depicting regions of IRF7 with precent identity of each region to Pa.IRF7 ortholog indicated below. (B) Sequence alignment of serine-rich domain at C terminus of Hs.IRF7 and Pa.IRF7. Conserved serines are indicated in green, and unique Pa.IRF7 serines in red. Orange and yellow boxes indicate residues selected for site-directed mutagenesis. (B) *STAT1*^−/−^ fibroblasts expressing IRF7 constructs indicated in (A) were infected with YFV at an MOI of 1. Cells were harvested for flow cytometry at 24h and infection was normalized to control cells. Modifications to the WT IRF7 protein sequence are indicated in bold on the left. Data are presented as means +/− SD, n=3. Statistical significance was determined by one-way ANOVA with Dunnett’s multiple comparison’s test. ****, P<0.0001, ns, not significant. (D) Representative western blot from one replicate of (C) showing equal IRF7 expression, differential IFIT1 induction, and actin loading control. (E) PaKi cells expressing indicated IRF7s were infected with ONNV at an MOI of 2 for 18h and infectivity was quantified by flow cytometry and presented as means +/− SD, n=3. Statistical significance was determined by one-way ANOVA with Sidak’s multiple comparison’s test. *, P<0.05; **, P<0.01; ns, not significant.

To explore the role of the serine-rich domain (SRD) in the STAT1-independent antiviral function of Pa.IRF7, we engineered several mutations, replacing certain serines with alanines or swapping bat and human residues to mimic each other’s sequence. The mutant IRF7s were expressed in human *STAT1*^−/−^ fibroblasts, which were then infected with YFV. All Ser-to-Ala mutations in Pa.IRF7 resulted in partial loss of activity (**Fig 6C**). Notably, the double mutant Pa.IRF7-S477A/S479A lost most of its antiviral activity, and its inhibitory effects were comparable to the modest effects seen with unmutated Hs.IRF7, which has serines at residues 477 and 479. Serine substitutions in Pa.IRF7-S480L/S485A double mutant also resulted in partial loss of antiviral activity. These data suggest important roles for multiple serines in Pa.IRF7-mediated antiviral function.

We then tried to convert Hs.IRF7 to a more Pa.IRF7-like antiviral protein. A Hs.IRF7-L480S/A485S double mutant, which contains 11 serines, as compared to 12 in Pa.IRF7 and 9 in Hs.IRF7, was not sufficient to increase the antiviral activity of Hs.IRF7. However, substituting L480 and A485 with phosphomimetic aspartic acids fully restored the antiviral function to levels seen with Pa.IRF7 (**Fig 6C**). Although these data are only correlative and do not pinpoint definitive phospho-sites, they suggest that the additional serines 480 and 485 in Pa.IRF7 may be important for conferring IFN-independent antiviral effects.

To understand the role of the serine-rich domain (SRD) in the antiviral effects of Pa.IRF7 within a bat cell context, we replaced the entire SRD of each IRF7 with its corresponding orthologous sequence. As a result, the Hs.IRF7(Pa.SRD) chimera incorporates all 12 serines from Pa.IRF7, in addition to the following amino acid substitutions: Val460Ala, His461Tyr, Thr465Val, Leu478Phe, Cys481Ala, Asp490Glu. In contrast, the Pa.IRF7(Hs.SRD) chimera contains all 9 serines from Hs.IRF7, along with the reciprocal amino acid substitutions mentioned above. Chimeric IRF7s were then expressed in PaKi cells, which were subsequently challenged with ONNV. Consistent with previous results, Pa.IRF7 strongly suppressed infection, whereas Hs.IRF7 was only minimally antiviral. Remarkably, Hs.IRF7-PaSRD significantly suppressed ONNV, similar to the efficacy of Pa.IRF7. This result demonstrates that the SRD from the bat protein is sufficient to confer Pa.IRF7-like antiviral effects to Hs.IRF7 in bat cells. Unexpectedly, Pa.IRF7 bearing the SRD from Hs.IRF7 (Pa.IRF7-HsSRD) also exhibited strong inhibition of ONNV, similar to native Pa.IRF7. Combined with the serine mutagenesis studies above, this outcome suggests that the antiviral potential of Pa.IRF7 may depend not only on the specific amino acid composition of the serine-rich domain, but also on how this region interfaces with the rest of the protein, which also has substantial divergence from Hs.IRF7 in certain domains.

## Discussion

In this study, we compared the antiviral activity of human and bat ISG orthologs and identified Pa.IRF7 as a strong inhibitor of several diverse RNA viruses. In contrast to ectopically expressed Hs.IRF7, which only has minor antiviral activity upon loss of IFN signaling, ectopically expressed Pa.IRF7 induces a robust ISG transcriptional response in human cells, even in the absence of a viral stimulus. Pa.IRF7 also contributes directly to ISG induction in bat cells, as demonstrated by CRISPR-mediated knockout studies. In mechanistic studies, we found that Pa.IRF7 has greater propensity than Hs.IRF7 to localize to the nucleus and directly bind ISG promoters. Domain swapping analysis and site-directed mutagenesis studies indicate that the serine-rich domain at the C terminus of Pa.IRF7 may contribute to the augmented antiviral phenotype.

Two previous studies have also examined unique properties of IRF7 from the black flying fox. Zhou et al. provided evidence that Pa.IRF7, despite unique sequence adaptations in a MyD88 binding region, was still able to induce IFN like other mammalian IRF7s (Zhou et al., 2014). While conducting our studies, Irving et al. demonstrated that elevated baseline expression of IRF1, IRF3, and IRF7 across various bat tissues enhances the basal antiviral state and contributes to the regulation of type I IFN ligands and ISGs (Irving et al., 2020). Their work revealed distinct subsets of genes regulated by each IRF, with IRF1 and IRF7 playing critical roles in inducing antiviral genes in the absence of IFN signaling. Irving et al. also provided evidence that bat IRFs, particularly Pa.IRF7, regulate a broad, IFN-like antiviral signature that is both basal and inducible in response to viral mimetics like dsRNA. Our studies complement the work of Irving et al. by showing similar IFN-independent antiviral activity of Pa.IRF7. We additionally provide mechanistic insight to this phenotype by showing that Pa.IRF7 directly binds ISG promoters and has additional C-terminal serines that appear to contribute to augmented antiviral activity.

The collective data on IRF7 from the black flying fox supports a model in which Pa.IRF7 induces early antiviral responses independently of traditional IFN signaling pathways, in contrast to the primary role of human IRF7 in amplifying IFN responses by directly inducing IFN gene expression. Importantly, we note that Hs.IRF7 has been shown to exhibit promiscuous transcriptional activity that activates IFNs and ISG expression in certain contexts. In the classical mode of Hs.IRF7 activation, kinases IKKε and TBK1 phosphorylate Hs.IRF7, leading to its activation and induction of IFN gene transcription (Fitzgerald et al., 2003; Sharma et al., 2003). This requirement for kinase-mediated activation underscores a regulated, conditional approach to initiating antiviral defenses, wherein Hs.IRF7 activity is tightly controlled and dependent on upstream signaling events. Notably, Lin et al. showed that overexpression of Hs.IRF7 stimulated IFN gene expression in the absence of viral stimulus (Lin, Mamane, et al., 2000). Schmid et al found that in the presence of ectopically expressed IKKε, Hs.IRF7 not only becomes activated but also gains the capacity to induce numerous ISGs (Schmid et al., 2010). This indicates that under specific conditions, Hs.IRF7 can directly contribute to the transcriptional activation of antiviral genes. This capacity for direct ISG induction reflects a more nuanced role for Hs.IRF7 in human antiviral immunity, capable of both amplifying IFN responses and inducing ISGs in a context-dependent manner. With respect to ISG induction, Pa.IRF7 appears to be distinct from Hs.IRF7 in that it can stimulate ISGs without a stimulus and without overexpression of IKKε, which we did not test in our studies.

The contrast between bat and human IRF7 underscores potential evolutionary adaptations in antiviral responses. While human IRF7 predominantly functions to amplify IFN responses through a well-defined pathway requiring upstream activation, our results suggest that Pa.IRF7 is pre-primed for a direct antiviral response, independent of such signaling cues. The additional serines at the C terminus of Pa.IRF7, absent in Hs.IRF7, may partly underlie this distinct functional capacity.

Our analysis presents several considerations to further our understanding of the divergent antiviral responses between Pa.IRF7 and Hs.IRF7. We hypothesize that Pa.IRF7 may have an inherently lower threshold for activation than Hs.IRF7, thereby contributing to enhanced nuclear localization and ISG promoter affinity. A pivotal outstanding question is whether the potent antiviral activity of Pa.IRF7 arises from its ability to be phosphorylated by basally expressed kinases, such as IKKε, TBK1, or some unknown kinase, in the absence of viral stimuli. The presence of additional unique serines in Pa.IRF7 may further augment its phosphorylation capacity or confer structural adaptations that enable easier kinase access. For example, the inhibitory domain may have increased plasticity and less autoinhibition, thus keeping Pa.IRF7 in a more constitutively active or readily activatable state. Drawing parallels with IRF3, where phosphorylation induces a conformational change to expose the transactivation domain (Lin et al., 1999; Qin et al., 2003), a similar activation mechanism could be at play for IRF7. A variability in activation thresholds, potentially lower in Pa.IRF7 due to its additional serines, may suggest an inherent structural predisposition for activation. Additionally, while differences in DNA affinity between IRF3 and IRF7 seem to be due to differences in their N-terminal DNA-binding domains (De Ioannes et al., 2011), differences between bat and human IRF7 may be more likely attributed to differences in the threshold for activation. Yet, the specific kinases responsible for this activation in Pa.IRF7, or whether a kinase is even necessary, require further study.

The hypothesis that Pa.IRF7 has constitutive or readily inducible activity suggests an evolutionary adaptation for an enhanced baseline antiviral defense, possibly offering preemptive protection against IFN pathway-targeting viruses. This unique antiviral strategy in bats may provide insights into novel approaches for modulating antiviral responses in humans. Future research should aim to identify the pathways and mechanisms responsible for Pa.IRF7 activation, determine structural differences underpinning its enhanced antiviral function, and understand the broader implications of these findings for bat physiology and potential applications in enhancing human antiviral defense mechanisms. Continued effort in these research areas will help provide a more comprehensive understanding of the innate immune responses across species.

## Methods

### Cell Lines

All cells were grown at 37°C, 5% CO_2_ and supplemented with 10% FBS (Life Technologies) and 1x nonessential amino acids (NEAA) (Life Technologies), except where noted. *STAT1*^−/−^ human fibroblasts (Dupuis et al., 2003), provided by J.-L. Casanova, were passaged in RPMI (Life Technologies). 293T and HeLa cells were passaged in DMEM (Life Technologies). BHK-21J, provided by C. Rice, were passaged in MEM (Life Technologies). THP-1 cells were grown in suspension with RPMI. PaKi cells (Crameri et al., 2009) were passaged in DMEM/F12 (Life Technologies) supplemented with 10% FBS only. Stable cells were maintained in 0.5 µg/ml (*STAT1*^−/−^ human fibroblasts) or 0.25 µg/ml (HeLa) of Puromycin.

### Viruses

YFV-Venus (derived from YF17D-5C25Venus2AUbi, gift from C. Rice), VEEV-GFP (derived from pTC83-GFP, gift from I. Frolov), ONNV-GFP (derived from pONNV.GFP, gift from S. Higgs), SINV-A-GFP (derived from pS300-GFP, gift from M, Heise), VSV-GFP (gift from J. Rose), and PIV3-GFP (based on strain JS, gift from P.Collins) were generated as previously described (Schoggins et al., 2014; Schoggins et al., 2011). For all viruses, virus-containing supernatant was centrifuged to remove cellular debris and stored at −80C until use. MOIs were derived from titering assays performed in *STAT1*^−/−^ fibroblasts (YFV) or Huh7.5 (VEEV, ONNV, SINV, VSV, PIV3) cells.

### Viral Infections

Cells were seeded into 24-well plates at a density of 8×10^4^ to 1×10^5^ cells per well, or from a 1:3 split if previously transduced. Viral stocks were diluted into media supplemented with 1% FBS to make infection media. Media was aspirated and replaced with 200 µL of infection media. Infections were carried out at 37°C for 1h, then 800-1500 µL media supplemented with 10% FBS was added back to each well. Cells were harvested for flow cytometry at the following times: ONNV: 18 h, SINV: 10 or 18 h, VEEV: 5-6 h, VSV: 4 h, YFV: 24 h, PIV3: 18 h.

### Lentiviral Pseudoparticle Production and Transduction

All lentiviral pseudoparticles were generated by co-transfecting 400,000 293T cells with expression plasmids pTRIP.CMV.IVSB.ISG.ires.TagRFP(Schoggins et al., 2011), pSCRPSY(Kane et al., 2016), HIV-1 gag-pol, and VSV-glycoprotein at a ratio of 5:4:1 (pTRIP) or 25:5:1 (pSCRPSY) as previously described. Cells were seeded at a density of 1-4×10^5^ cells per well on 6 or 24-well plates. The following day, lentiviral psuedoparticles diluted in PP media (media supplemented with 3% FBS, 1x NEAA, and 4 µg/mL polybrene) were added to cells and cells were spinoculated at 800 x *g*, 45 min, 37°C. Following spinoculation, media was changed to fresh media containing 10% FBS and 1x NEAA. Stable cells were selected by Puromycin and scaled up accordingly.

### Flow cytometry

Cells were detached using Accumax (Innovative Cell Technologies), fixed in 1% PFA for 10 min at room temperature, and pelleted by centrifugation at 800xg. Fixed cell pellets were resuspended in 200 µL FACS buffer (1X PBS supplemented with 3% FBS). Samples were run on a Stratedigm S1000 instrument using CellCapTure software and gated based on live cells, singlets, RFP signal for ISG expression, and finally GFP signal for viral infection. Data analysis was done using FlowJo software.

### ISG Screens

*STAT1^−/−^* human fibroblasts ere seeded at a density of 1×10^5^ cells per well on 24-well plates. The following day, cells were transduced with pTRIP.CMV.IVSb.ires.TagRFP lentiviral pseudoparticles, leading to expression of an ISG of interest and RFP in a one-well: one-ISG format. After 48h, cells were split 1:3. The following day, cells were infected with a GFP-reporter virus and harvested for flow cytometry as described above.

### Western Blots

Samples for WB were prepared one of two ways. Cells were detached from plates via Accumax, pelleted, and washed with PBS, followed by addition of RIPA buffer (150 mM NaCl, 1% Triton X-100, 0.5% Sodium deoxycholate, 50 mM Tris pH 8.0, 0.1% SDS, 1 Protease tablet). Samples were incubated on ice for 15 minutes, followed by shearing with syringe/needle, and centrifugation. Lysate was transferred to fresh microcentrifuge tube. Protein concentration was determined by Pierce BCA Protein Assay Kit (Fisher). SDS loading buffer (0.2M Tris-Cl pH6.8, 5% SDS, 25% glycerol, 0.025% Bromophenol Blue, 6.25% 2-ME) was added to equal protein concentrations for a final concentration of 1X. Alternatively, samples were prepared by aspirating media from wells, washing with PBS, and lysing directly in the well with 1X SDS loading buffer (0.04M Tris-HCl pH 6.8, 2% SDS, 2mM BME, 4% glycerol, 0.01% bromophenol blue). All samples were boiled for 5 min and immediately frozen and kept at −20°C until use. Samples were loaded on a 12% polyacrylamide gel (TGX FastCast Acrylamide Kit, Bio-Rad), and run at 100-250V for 30-90 min. Polyacrylamide gels were transferred to PVDF or nitrocellulose membranes using either wet transfer (100V, 45 min, 4°C) or semi-dry transfer (7 min, RT). Membranes were blocked with 5% nonfat milk in TBST (50 mM Tris-HCl pH 7.4, 150 mM NaCl, 0.1% Tween-20) for 1h to overnight, then incubated with primary antibody 1-2h, washed 3x with TBST for 5 min each, incubated in secondary antibody for 30 min, washed 3-5x with TBST, and finally developed using ECL Western Blotting Substrate (Thermo Scientific Pierce) or imaged on Licor.

All primary and secondary antibodies were diluted in TBST. Primary antibodies include anti-IRF7 (Abclonal A0159), anti-IRF7 (Cell signaling 4920), anti-Flag M2 (Sigma F3165), anti-GAPDH (Proteintech 60004-1), anti-STAT1 (Proteintech 66545-1), anti-IFIT1 (Proteintech 23247-1), anti-LaminB (Proteintech 66095-1), and anti-beta Actin (Abcam ab6276). Secondary antibodies include Goat anti-Mouse IgG (H+L) HRP (Life Technologies), Goat anti-Rabbit IgG (H+L) HRP (Life Technologies), Donkey anti-Mouse IgG (H+L) highly cross-adsorbed HRP (Life Technologies), Donkey anti-Rabbit IgG (H+L) highly cross-adsorbed (Life Technologies), IRDye 800CW Goat anti-Rabbit IgG (Licor), and IRDye 680RD Goat anti-Mouse IgG (Licor).

### Microarray

*STAT1*^−/−^ fibroblasts stably expressing IRF7 or vector control were seeded in 6-well plates at a density of 4×10^6^. The following day, RNA was harvested using the RNeasy Mini Kit (Qiagen) following the manufacturer’s instructions and stored at −80°C until use. The RNA integrity score (RIN) was 10 for all samples. Samples were run on an Illumina Human HT 12 v4 chip. Microarray data have been deposited at NCBI GEO with accession number GSE265790.

### IRF7 KO in PaKi cell line

pSpCas9(BB)-2A-GFP (PX458) was obtained from F. Zhang (Addgene plasmid # 48138; http://n2t.net/addgene:48138; RRID:Addgene_48138). Two sgRNA sequences were cloned into pSpCas9(BB)-2A-GFP following previously published protocol (Ran et al., 2013). PaKi cells were plated at a density of 250,000 cells per well in 6-well format. 1500 ng each sgRNA was co-transfected in each well using Lipofectamine 3000 (Life Technologies). Cells were incubated for 2-3 days at 37°C with 5% CO2. Cells were sorted for GFP expression and plated as single cell clones. Western Blots were performed on individual clones to determine loss of IRF7. Genomic DNA was isolated from IRF7 KO clone 4G9, and a fragment containing location of sgRNA was amplified by PCR. Fragment was sent for sequencing to determine gene edit.

### IRF7-3xFLAG endogenous tagging in PaKi cell line

PaKi cells were seeded at a density of 2×10^6^ cells in 10 cm plates. The following day, cells were co-transfected with 1ug lentiCRISPRv2 and 4 ug donor homology plasmid using Lipofectamine 3000 according to the manufacturer’s protocol. The lentiCRISPRv2 plasmid encodes a guide RNA that targets 6 nucleotides upstream of the stop codon. After 4h, media was changed for fresh media containing 0.1uM SCR-7 (Tocris) to promote homology directed-repair. At 48h after transduction, cells were selected with 4ug/mL of puromycin for 72h. The surviving bulk population was tested for an IFN-inducible FLAG signal by western blot. Single-cell clones were generated by limiting dilution and screened for IRF7-3xF expression by WB. Genetic confirmation of the 3xFLAG tag was done by genomic PCR and TOPO cloning, indicating that all alleles has been modified.

### DNA Constructs and Plasmid Propagation

The human ISG library has been previously described (Schoggins et al., 2011). The bat ISG library was created by subcloning codon-optimized ISG ORFs into a pENTR221 backbone purchased from GeneCopoeia. The ORFs were moved into pTRIP.CMV.IVSb.ires.TagRFP-DEST using LR Clonase II (Invitrogen) according to manufacturer’s instructions. Correct assembly was verified by Sanger sequencing. pTRIP.CMV.IVSb.ires.TagRFP.Fluc (firefly luciferase) was used as a negative control in the screens. For Pa.IRF7, the original library contained an IRF7 sequence corresponding to accession number ELK13169.1. This older accession encoded a truncated protein that lacked 30 amino acids relative to the updated full length Pa.IRF7 with accession XP_006911017.1.

Plasmids for stable expression pSCRPSY.B-IRF7, pSCRPSY.H-IRF7 were generated by Gateway cloning of pENTR vectors into pSCRPSY using LR Clonase II according to the manufacturer’s instructions. Correct assembly was confirmed by Sanger sequencing. pSCRPSY.Empty is used as a vector control.

To generate single and double-point IRF7 mutants, pENTR221 vectors containing open reading frames for bat or human full-length IRF7 (pENTR221.H-IRF7.3XFL and pENTR221.B-IRF7.3XFL) were digested with BamHI and EcoRV or BgII and BamHI, respectively. The large fragment was purified using the QIAquick Gel Extraction Kit (Qiagen). Point mutations were added by Gibson (NEB) assembly of the large fragment and a small synthesized (Genewiz) fragment containing the mutation of interest following the manufacturer’s directions. To generate serine rich domain swaps, pENTR221 vectors (pENTR221.H.IRF7.3XFL and pENTR221.B-IRF7.3XFL) were digested with BamHI and EcoRV or BgII and ClaI, respectively. The large fragment was purified and a gBlock (IDT) fragment containing serine swaps was ligated using Gibson assembly. Correct assembly was verified using Sanger sequencing. The ORFs were then subcloned via Gateway technology into pTRIP.CMV.IVSb.ires.TagRFP or pSCRPSY for use as expression vectors and correct assembly confirmed by Sanger sequencing.

### qRT-PCR

Total RNA was isolated using RNeasy Mini Kit per manufacturer protocol (Qiagen). Reactions were prepared with the QuantiFast SYBR Green RT-PCR kit (Qiagen), using 50 ng total RNA per reaction. Samples were run on the Applied Biosystems 7500 Fast Real-Time PCR System or QuantStudio 3 Real-Time PCR System. QuantiTect Primer Assays (Qiagen) were used to amplify IFI44L (QT00051457), RSAD2 (QT00005271), and IFNB1 (QT00203763). Primers used to amplify human genes: OAS2 5’-GAACACCATCTGTGACGTCCT-3’ and 5’-GAGCCACCTATGGCCACTCC-3’, RPS11 5’-ATCCGCCGAGACTATCTCCA-3’ and 5’-GGACATCTCTGAAGCAGGGT-3’. Primers used to amplify bat genes: IRF7 5’-GGCTGGAAAACCAACTTCCG-3’ and 5’-CTCCCCCTGGTTAATGCCTG-3’, OAS1 5’-TGAAGCAGAGACCAGCCAAG-3’ and 5’-TTTCCACGTTCCCAAGCGTA-3’, USP18 5’-TCGGCAGATCCTGTTGAGAAG-3’ and 5’-TGTTGTGTAAACCGACTGGG-3’, RPS11 5’-ATCCGCCGAGACTATCTCCA-3’ and 5’-GGACATCTCTGAAGCAGGGT-3’. Relative expression was calculated using 2^−ΔΔCT^ method.

### Subcellular Fractionation Assay

2.5×10^6^ THP-1 or PaKi cells were treated with 100 U Universal IFN (PBL Assay Science) for 18 hrs at 37°C and 5% CO_2_. At 18 hrs, THP-1 cells were collected, pelleted, and washed 1x DPBS. Paki cells were dislodged with Accumax, pelleted, and washed 1x with DPBS. Subcellular fractionations were prepared using Cell Fractionation Kit (Cell Signaling) per manufacturer protocol, with the following exception. Benzonase was added to CyNIB buffer. 1 µl Benzonase was added to whole cell lysate fraction, incubated on ice for 30 min and nuclear fraction was incubated at room temp for 30 min. Both samples were sheared with needle/syringe and centrifuged at 7800 xg for 10 min at 4°C. Lysate was transferred to new microcentrifuge tube and 5X SDS-SB was added to a final concentration of 1X. 5X SDS-SB was added to cytosolic fraction to a final concentration of 1X. All samples were boiled at 95°C for 5 min. Protein was loaded on 12% acrylamide gel, transferred to PVDF or nitrocellulose membrane, followed by Western Blotting.

### ISRE pulldown assays

Lysates from *STAT1^−/−^* fibroblasts stably expressing vector control or bat or human IRF7 were prepared the same way as for the dimerization assays. 50 μL lysate (∼25 ug total protein) was mixed with 2 μg 5’-biotinylated dsDNA probe and 50 μL streptavidin agarose resin (Thermo Scientific) in PBS containing protease inhibitors (Roche) and phosphatase inhibitors (Roche) to a final volume of 250 μL. The mixture was gently rotated at 4°C overnight.

## ACKNOWLEDGEMENTS

This work was funded in part by grants to J.W.S: NIH Grants AI117922 and AI158124, The Welch Foundation (I-2013-20220331), and the UT Southwestern High Impact / High Risk Grant. John W. Schoggins holds an Investigators in the Pathogenesis of Infectious Disease Award from the Burroughs Wellcome Fund. We thank Ian Boys for unpublished evolutionary analyses on IRF7 and Dustin Hancks for helpful discussions.

---

**Figure S1.**
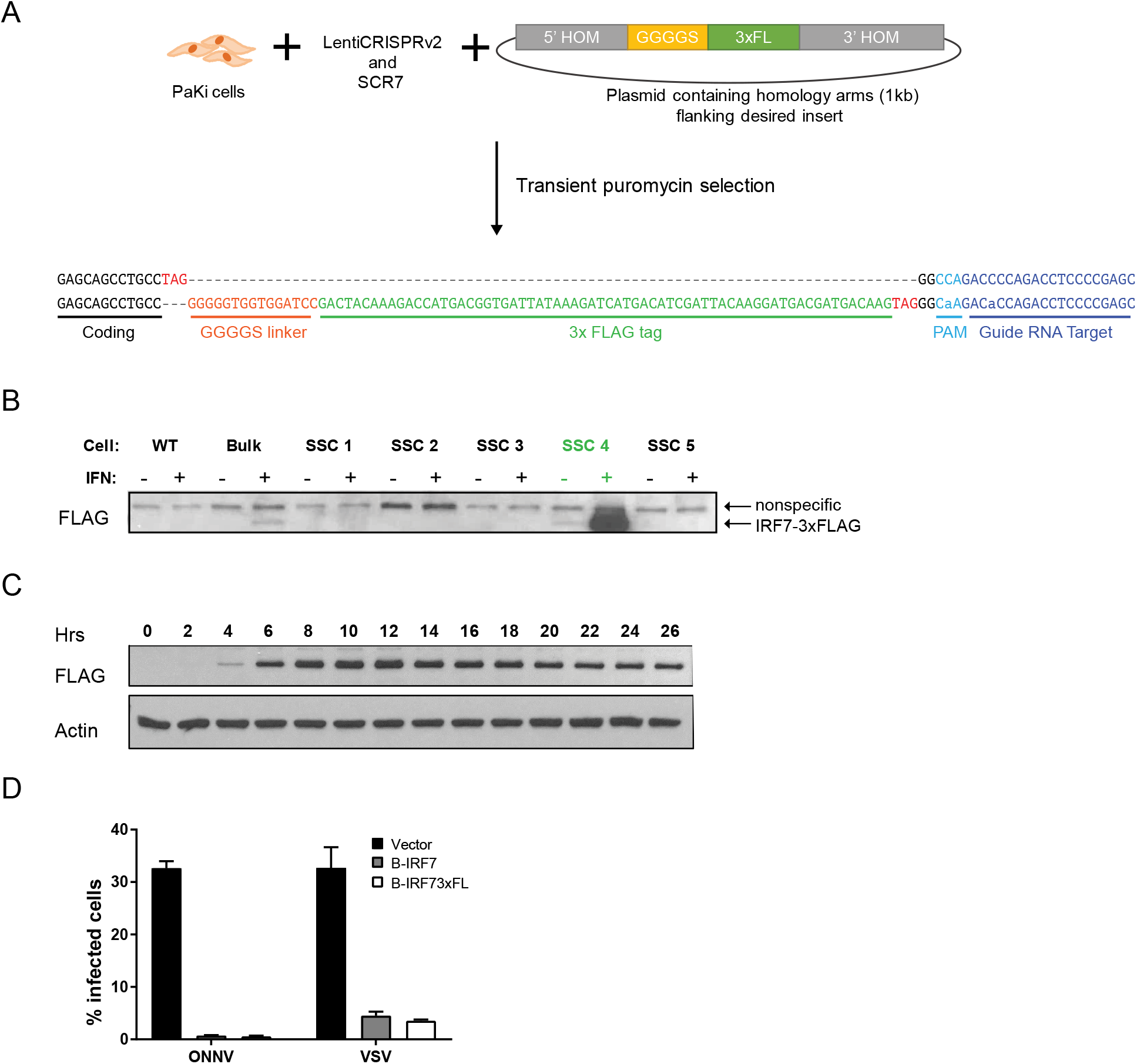
Strategy to tag endogenous Pa.IRF7 with 3X-FLAG epitope using homology-directed repair. (A) PaKi cells were transfected with lentiCRISPRv2 and gRNAs targeting the C-terminal region of IRF7 along with a donor plasmid containing a linker and 3xFLAG tag flanked by ∼500bp upstream and downstream of the CRISPR target site. SCR7 was added to promote homology-directed repair. Cells were selected with puromycin for 3 days and then expanded for analysis. (B) Western blot of untagged (WT) PaKi cells, bulk transfected cells, and single cell clones (SSC), without and with IFN treatment. Bulk cells and SSC4 have an IFN-inducible FLAG signal. (C) PaKi IRF7-3xFLAG cells (SSC4) were treated with 50U/mL of universal IFNα. Lysates were harvested at indicated times and analyzed by Western blot, probing with anti-FLAG antibody. (D) *STAT1*^−/−^ fibroblasts were transduced with lentivirus expressing Pa.IRF7 or Pa.IRF7 with a C-terminal 3xFLAG tag identical to that used in endogenously-tagged PaKi cells. Cells were then infected with ONNV or VSV at an MOI of 1 and infection was quantified via flow cytometry, n=2.

**Figure S2.**
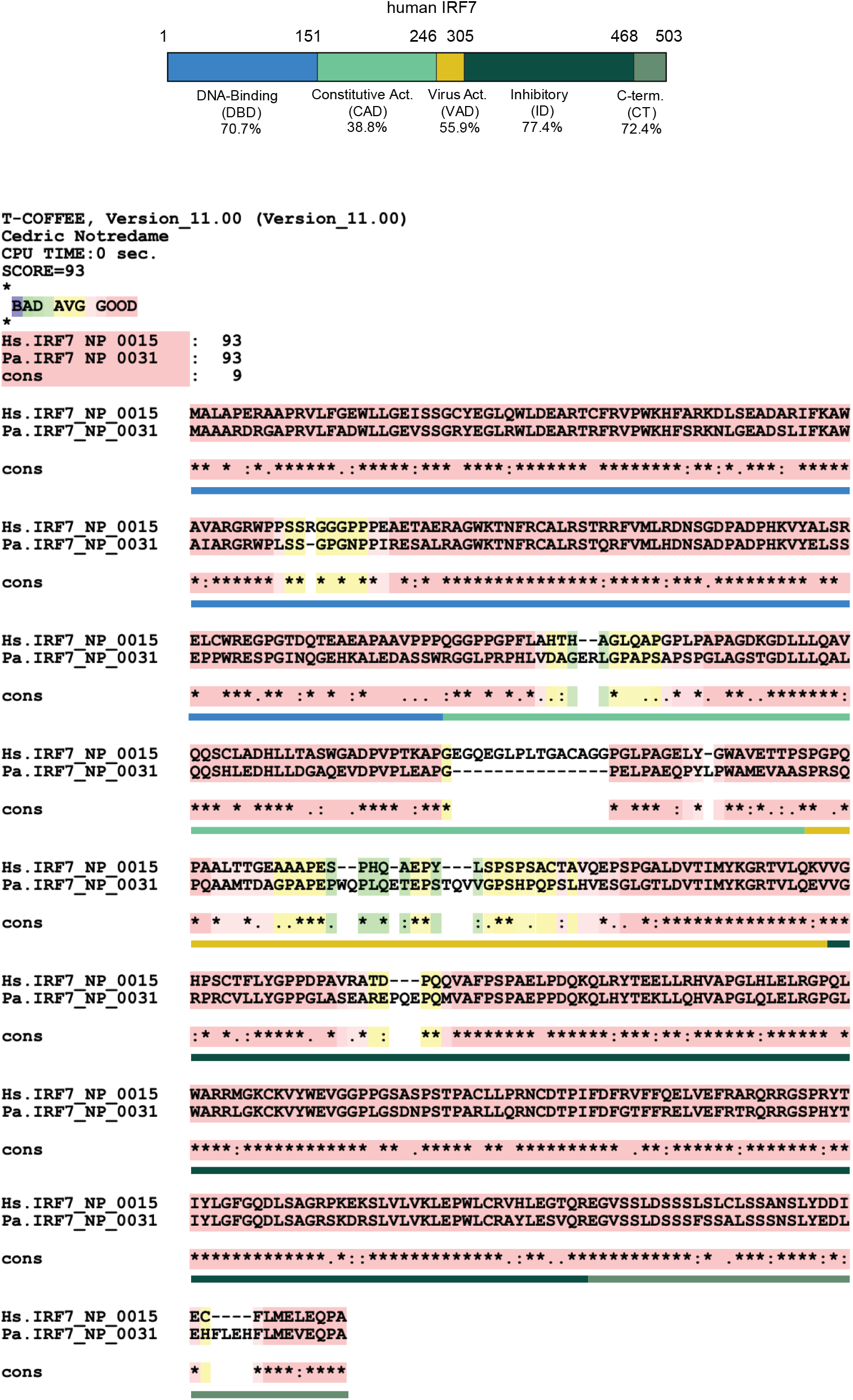
Alignment and homology of Hs.IRF7 and Pa.IRF7. (Top) Cartoon diagram of Hs.IRF7 depicting regions of IRF7 with precent identity of each region to Pa.IRF7 ortholog indicated below. (Bottom) T-COFFEE alignment of Hs.IRF7 and Pa.IRF7 proteins. Colored bars underneath alignment match colored regions in cartoon in (A).

**Table S1.**
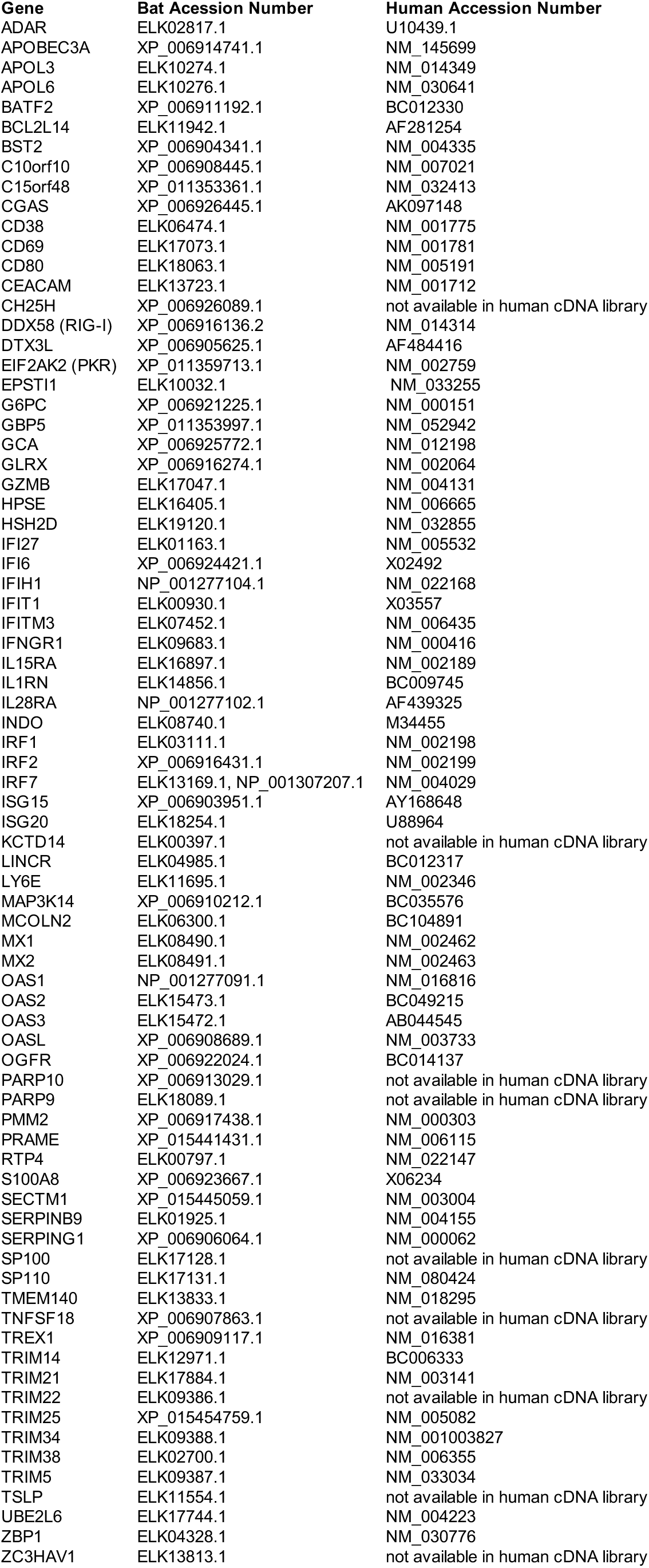
Accession numbers of bat and human ISGs for functional screens.

**Table S2.**
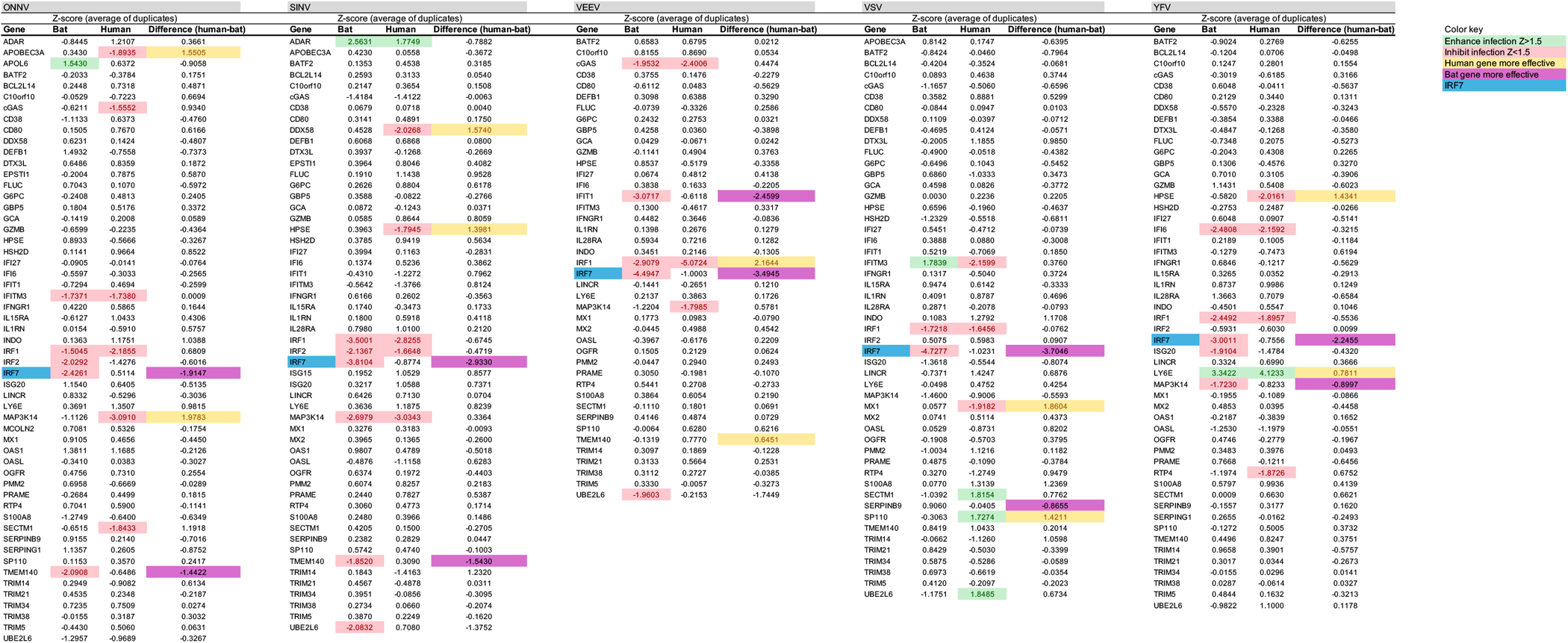
Z-scores from ISG screens used to generate Fig 1.

